# Protective Effects of Edaravone Against Hypoxia-Induced Lethality in Male Swiss Albino Mice

**DOI:** 10.1101/2020.05.22.111401

**Authors:** Fatemeh Shaki, Mina Mokhtaran, Maedeh Raei, Amir Shamshirian, Shahram Eslami, Danial Shamshirian, Mohammad Ali Ebrahimzadeh

## Abstract

Edaravone has been recently used for the treatment of acute cerebral infarction. However, there is no data on the protective effects of this drug against hypoxia-induced lethality. Here, we aimed to evaluate the protective effects of edaravone against hypoxia-induced lethality and oxidative stress in mice through three experimental models of hypoxia.

**Materials and Methods:** male Swiss albino mice were randomly housed in groups of 10. Mice received single i.p. injections of edaravone for four consecutive days. Three experimental models of hypoxia: asphyctic, circulatory, and haemic were applied in this study. Oxidative stress, lipid peroxidation and glutathione content were assessed after hypoxia induction.

**Results:** Significant protective activities were observed in all models of hypoxia. Antihypoxic activities were highly significant in asphytic and circulatory hypoxia. These effects were dose-dependent. Edaravone, at 5 mg kg^−1^, showed statistically significant activities concerning the control groups. Edaravone significantly prolonged the latency for death. At 2.5 mg kg^−1^, it prolonged survival time (26.08 ± 0.79 min). This effect was statistically significant (*P*<0.05). Also, edaravone significantly inhibited hypoxia-induced oxidative stress (lipid peroxidation and glutathione oxidation) in three models of hypoxia.

**Conclusions:** Edaravone showed an excellent protective effect against hypoxia in all tested models and decreased the oxidative stress in the brain tissue of hypoxic mice. Notably, results showed significant and dose-dependent effects on the models of asphytic and circulatory hypoxia. The antioxidant activity itself might be a proposed mechanism for the antihypoxic activities of this drug.

**Impact statement:** This study provides a novel proof-of-concept for a FDA approved drug, which will open a new area of research in the field and drug repurposing.

## Introduction

The imbalance between the oxygen supply and its demand regulates organ hypoxia. It occurs mainly in ischemia and heart diseases, leading to numerous harmful effects in different tissues, especially in the brain ^1^. The brain has a high oxygen usage, which comprises 20-25% of total body O_2_ consumption; therefore, it is highly susceptible to hypoxia.

Edaravone, a novel and potent free radical scavenger, has been shown to have protective effects against cerebral ischemia-reperfusion injuries in some experimental animal models^2^. It has been reported that edaravone can inhibit the activation of lipoxygenase and peroxidation of the phosphatidylcholine liposomal membrane *in vitro*^3^. The clinical efficacy of edaravone against ischemic brain attack has been demonstrated by the presence of significant improvements in functional outcomes in a human study^3, 4^. Systemic administration of edaravone attenuated the increase of malondialdehyde levels, reduced superoxide dismutase activity, and suppressed retinal dysfunction after retinal ischemia/reperfusion in rats^5^.

However, as far as we know, there were no data available on the effects of edaravone in hypoxic conditions. The present study aimed to determine the antihypoxic activities of edaravone to understand a possible mechanism of its action in cerebrovascular diseases.

## Material and methods

### Experimental animals and diet

Male Swiss albino mice (20 ± 2 g) were randomly housed in groups of 10 in polypropylene cages at ambient temperature, 25 ± 1°C and 45-55% relative humidity, with a 12 h light: 12 h dark cycle. The animals had free access to standard pellet and water *ad libitum*. All the experimental procedures were conducted following the NIH guidelines of the Laboratory Animal Care and Use. The Animal Ethics Committee of Mazandaran University of Medical Sciences approved the experimental protocol due to the code IR.MAZUMS.REC.93.976.

### Hypoxia models

Three experimental models of hypoxia were applied in this study. In each model, sixty mice were divided into ten groups of six mice. The asphyctic hypoxia model induced the animals by putting them individually in a tightly closed 300 ml glass container placed underwater in an aquarium of 25°C. Mice received single i.p. injections of 2.5 and 5 mg kg^−1^ doses of edaravone for four consecutive days. On the 4^th^ day, edaravone or phenytoin (50 mg kg^−1^) was injected 30 min before they were subjected to hypoxia. Circulatory and Haemic hypoxia models were applied to the animals through i.p. injection of NaF (150 mg/kg) and NaNO_2_ (360 mg/kg), respectively, on the 4^th^ day, 30 minutes after *i.p.* administration of edaravone. In both models, mice received single i.p. injections of 2.5, 5 or 10 mg kg^−1^ doses of edaravone for four consecutive days. Finally, the death latencies were recorded for all models. Negative control groups were treated with normal saline and propranolol used as the positive control^6^.

### Oxidative stress assessment

Sixty mice were divided into ten groups of six mice. Mice were exposed to hypoxia by the above three methods. After a cut off time, the animals were anesthetized by ketamine (80 mg/kg) and xylazine (5 mg/kg). Consequently, the brain tissue was separated, minced and homogenized with a glass handheld homogenizer. Then, tissue homogenates were centrifuged at 3000xg for 10 minutes at 4℃ and supernatant used to assess oxidative stress markers, including lipid peroxidation (LPO) and Glutathione (GSH). The content of malondialdehyde (MDA) was determined. The MDA amount was assessed by measuring the absorbance at 532 nm with an ELISA reader (ELX800, Biotek, USA). Tetramethoxypropane was used as standard, and MDA content was expressed as nmol/mg protein. GSH content was determined by DTNB as an indicator and read at 412 nm on a spectrophotometer (CE2501, CECIL, France) and expressed as μM^7^.

### Statistical Analysis

Data were presented as Mean ± SEM. Analysis of variance (ANOVA) was performed. Duncan’s new multiple-range test was used to determine the differences in means. All *p* values less than 0.05 were considered statistically significant. Statistical analyses were performed with GraphPad Prism v.6 (GraphPad Software Inc, San Diego, CA).

## Results

The results of asphytic hypoxia are shown in Figure 1a. The results showed that the effect was dose-dependent. At 2.5 mg kg-1, edaravone showed statistically significant activity concerning the control group (*P*<0.05). At 5 mg kg^−1^, it significantly prolonged the latency for death concerning the control group (33.10 ± 1.99 vs. 21.25 ± 0.63 min, *P*<0.0001).

**Figure 1Fig. 1.**
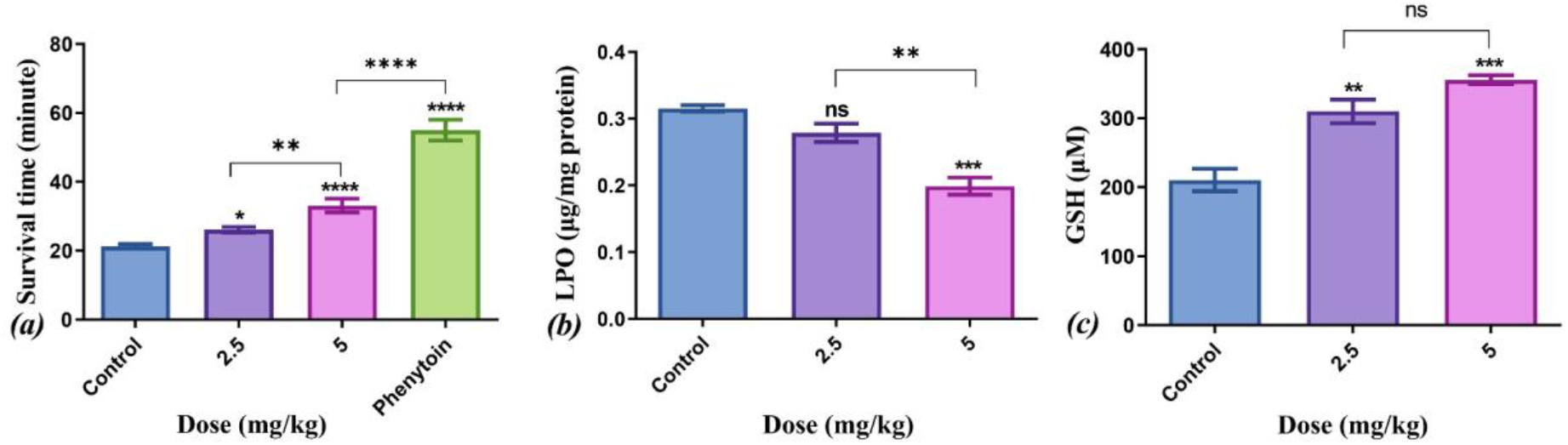
a: Antihypoxic activities of edaravone in asphyctic hypoxia in mice. b. Effects of Edaravone on asphyctic hypoxia-induced lipid peroxidation in brain tissue in mice. c. Effects of Edaravone on asphyctic hypoxia-induced GSH oxidation in brain tissue.

The results of the circulatory hypoxia are shown in Table 1. Edaravone at 2.5 mg kg^−1^ dose did not show any activity but at 5 mg kg^−1^ dose was substantially effective. At this dose, it significantly prolonged the latency for death compared to the control group (28.30 ± 2.70 vs. 9.78 ± 0.41 min, *P*<0.0001). This effect was dose-dependent.

**Table 1.** Antihypoxic activities of edaravone in haemic and circulatory hypoxia in mice

| Dose (mg/kg) | Haemic hypoxia |  |  | Circulatory hypoxia |  |  |
| --- | --- | --- | --- | --- | --- | --- |
|  | Hypoxia | MDA | GSH | Hypoxia | MDA | GSH |
| NS | 8.01±0.29 | 0.273±0.003 | 211.3±6.90 | 9.78±0.41 | 0.222±0.007 | 183.0±7.83 |
| 2.5 | -- | -- | -- | 12.05±0.55<sup>ns</sup> | 0.214±0.006<sup>ns</sup> | 229.3±6.98<sup>***</sup> |
| 5 | 9.57±0.50<sup>ns</sup> | 0.274±0.008<sup>ns</sup> | 304.3±9.25<sup>****</sup> | 28.30±2.70<sup>****</sup> | 0.177±0.007<sup>****</sup> | 239.3±8.79<sup>****</sup> |
| 10 | 13.14±0.82<sup>****</sup> | 0.200±0.006<sup>****</sup> | 303.8±3.55<sup>****</sup> | -- | -- | -- |
| Pro. 20 | 16.44±0.62<sup>****</sup> | -- | -- | 12.76±0.40<sup>ns</sup> | -- | -- |
| Pro. 30 | -- | -- | -- | 16.16±0.30<sup>*</sup> | -- | -- |

MDA: Malondialdehyde (µg/mg protein); GSH: Glutathione content (µM); Pro.: Propranolol; Data were expressed as Mean ± SEM (n = 6). Ns: not significant; \*\*\* $P<0.001$ , \*\*\*\* $P<0.0001$

This compound showed promising activity in the haemic model (Table 1). The control group died because of hypoxia in 8.01 ± 0.29 min. Edaravone at 5 mg kg^−1^ prolonged the latency for death, but this activity was not statistically significant (9.57 ± 0.50 min, *P*>0.05). At 10 mg kg^−1^, edaravone showed statistically considerable activity compared to the control group (*P*<0.0001). It significantly prolonged the latency for death (13.14 ±0.82 min, *P*<0.0001).

Elevation of MDA is known as an important marker for oxidative stress. The level of MDA as an indicator of lipid peroxidation in the asphytic hypoxia model is shown in Figure 1b. Edaravone as a dose-dependent drug hugely prevented the asphytic hypoxia-induced LPO concerning the control. Even at 2.5 mg kg^−1^, this decrease was statistically significant (*P*<0.01). At a higher dose (i.e. 5 mg kg^−1^), this reduction was more effective (P< 0.0001). As shown in Table 1, edaravone at both doses decreased the lipid peroxidation due to circulatory hypoxia conditions (P<0.0001). Also, the assessment of MDA concentration in the haemic model showed that edaravone significantly (*P*<0.0001) inhibited the increased lipid peroxidation induced by haemic hypoxia at a dosage of 10 mg kg^−1^ (Table 1).

The GSH levels (as the main intracellular antioxidant) in the brain tissue of asphytic hypoxia mice treated with edaravone (2.5 and 5 mg kg^−1^) was significantly (*P*<0.0001) increased in comparison with the control group (Fig. 1c). At 2.5 mg kg^−1^, edaravone increases GSH level from 210.5 ± 14.04 to 310.1± 14.82 μM.

As shown in Table 1, edaravone at both doses of 2.5 and 5 mg kg^−1^ significantly inhibited GSH oxidation due to the circulatory hypoxia conditions (*P*<0.001 and P< 0.0001, respectively). The assessment of GSH concentration in the haemic model showed that edaravone significantly inhibited the increased lipid peroxidation induced by haemic hypoxia at both doses of 5 and 10 mg kg^−1^ (P< 0.0001).

## Discussion

Statistically significant antihypoxic activities were found in 5 mg kg^−1^ dose of edaravone in experimental models of hypoxia in mice. At this dose, edaravone showed statistically significant activity compared to the control group. This effect was dose-dependent, and at higher doses, this drug showed significantly higher effects. Protective effects of edaravone have been reported against cerebral ischemia-reperfusion injuries in some experimental animal models^3, 4^. A close association between oxidative metabolism and cholinergic function has been found during the investigations of NaNO_2_ on brain metabolism. Chemical hypoxia is induced by the NaNO_2_ injection, which decreases the oxygen-carrying capacity of the blood by converting hemoglobin to methemoglobin. This toxic dose is injected 30 min after the phenolic therapy. Immediately after the NaNO_2_ injection, animals are placed in small cages, and the time between injection of NaNO_2_ and respiration cessation is recorded. Edaravone showed moderate protective activity in the haemic model.

Available research studies illustrate that using NaF induces circulatory hypoxia, increasing blood histamine content and decreasing oxygen-carrying capacity. Edaravone at 5 mg kg^−1^ was highly effective in this model. The mechanism of this protective action might be due in part to the antioxidant activity of edaravone. Because there is no standard drug for haemic and circulatory hypoxic models, the results of this study were compared to those of control groups.

Assessment of oxidative stress markers in the brain tissue of mice in three hypoxia models showed that edaravone reduced the level of oxidative stress compared to the control group. The hypoxia-induced ROS production can cause oxidation of other cellular macromolecules such as protein, DNA, RNA, lipid peroxidation, neuronal dysfunction or death^8–10^. Lipid peroxidation is one of the most critical indicators of oxidative stress that has harmful effects on cells or tissues and can lead to more free radical production via a chain reaction, and finally, cell membrane disruption.

The present study revealed that hypoxia causes lipid peroxidation in brain tissue, which significantly inhibited by edaravone. Previous studies showed that edaravone prevented lipid peroxidation and nitric oxide production in the neonatal rat brain following hypoxic-ischemic insult and led to attenuation of neuronal damage in the neonatal rat brain^11, 12^. Another study demonstrated that the prophylactic administration of edaravone ameliorated transient hypoxic-ischemic brain injury *via* reducing oxidative stress^13^.

In cerebral hypoxia, phenytoin afforded significant protection of neurons in the hippocampus and the dentate nucleus compared to saline. Four mechanisms could contribute to this apparent protective effect of phenytoin. Phenytoin decreases the cerebral metabolic rate of oxygen consumption by 40-60%, decreases cerebral lactate production, and increases the cerebral levels of glucose, glycogen, and phosphocreatine, which can be expected to raise tolerance to brain ischemia. Under conditions of total brain ischemia, phenytoin has been shown to increase cerebral blood flow by vasodilation. Cerebral ischemia is associated with increases in extracellular potassium. Phenytoin inhibits this increase in K^+^, suggesting that this effect is part of its protective action by diminishing the edema following cerebral ischemia. Phenytoin stabilized neuronal membranes and prevented the rise in intracellular sodium, usually seen under cerebral hypoxic conditions due to the facilitation of the Na^+^-K^+^ ATPase system. The stabilizing action of phenytoin on membranes might decrease some of the morphologic and biochemical changes seen after cerebral hypoxia. Cerebral protective effects of phenytoin can be explained by blockade of Ca^2+^ channel, which will decrease intracellular Ca^2+^ that causes the inhibition of free fatty acid liberation toward the protection of the cellular membrane^14, 15^.

Catecholamines, particularly norepinephrine, are released into the blood when mammals are exposed to hypoxia or any number of other stressors and increase the sensitivity of heart muscle to lack of oxygen. Propranolol inhibits ventilatory response to norepinephrine in hypoxia^16^. It protects the fine ultrastructure of heart muscle against damage caused by hypoxia and protects mitochondrial function by the maintenance of near-normal mitochondrial oxidative phosphorylating and Ca^2+^ accumulating activities after hypoxic perfusion. Studies have shown that propranolol protects heart muscle against the effects of hypoxia and ischemia^17, 18^.

Glutathione (GSH) is the primary antioxidant in the cellular system that directly scavenges free radicals^19^. Previous studies showed a fall in GSH level. In the present study, after hypobaric hypoxia, that might be due to the inhibition of GSH synthesis and increased utilization of GSH for detoxification of hypoxia-induced free radical^20^. Further, a decreased GSH concentration has been observed in the present study, leading to an aggravation of oxidative stress levels and more brain injury. Treatment with edaravone significantly restored GSH concentration in brain tissue after hypoxia conditions.

## Conclusions

Edaravone showed an excellent protective effect against hypoxia in all tested models and decreased the oxidative stress in the brain tissue of hypoxic mice. Notably, results showed significant and dose-dependent effects on the models of asphytic and circulatory hypoxia. The antioxidant activity itself might be a proposed mechanism for the antihypoxic activities of this drug.

## Authors’ contributions

F.S., D.S, and MA.E. designed the study; F.S., M.M, M.R and S.E carried out the experiments; MA.E. and A.S. analyzed the data; A.S. wrote the manuscript in consultation with M.M, D.S. and MA.E.; and all of the authors read and revised the final version of the manuscript.

## Acknowledgments

The authors express their appreciation to the Vice-Chancellor for Research at Mazandaran University of Medical Sciences for supporting this experimental study.

## Declarations of competing interests

The authors declared no potential conflicts of interest with respect to the research, authorship, and publication of this article.

## Funding

This study funded by the Vice-Chancellor for Research at Mazandaran University of Medical Sciences [grant number 1393-976].

## Data Availability

The datasets generated and/or analyzed during the current study are available from the corresponding author on reasonable request.

## Notes

### Competing Interest Statement

The authors have declared no competing interest.

### Summary of Updates

Revised and it is under review at Experimental Biology and Medicine

